# Using networks to analyze and visualize the distribution of overlapping reading frames in virus genomes

**DOI:** 10.1101/2021.06.10.447953

**Authors:** Laura Muñoz-Baena, Art F. Y. Poon

## Abstract

Gene overlap occurs when two or more genes are encoded by the same nucleotides. This phenomenon is found in all taxonomic domains, but is particularly common in viruses, where it may increase the information content of compact genomes or influence the creation of new genes. Here we report a global comparative study of overlapping reading frames (OvRFs) of 12,609 virus reference genomes in the NCBI database. We retrieved metadata associated with all annotated reading frames in each genome record to calculate the number, length, and frameshift of OvRFs. Our results show that while the number of OvRFs increases with genome length, they tend to be shorter in longer genomes. The majority of overlaps involve +2 frameshifts, predominantly found in ds-DNA viruses. However, the longest overlaps involve no shift in reading frame (+0), increasing the selective burden of the same nucleotide positions within codons, instead of exposing additional sites to purifying selection. Next, we develop a new graph-based representation of the distribution of OvRFs among the reading frames of genomes in a given virus family. In the absence of an unambiguous partition of reading frames by homology at this taxonomic level, we used an alignment-free k-mer based approach to cluster protein coding sequences by similarity. We connect these clusters with two types of directed edges to indicate (1) that constituent reading frames are adjacent in one or more genomes, and (2) that the reading frames overlap. These adjacency graphs not only provide a natural visualization scheme, but also a novel statistical framework for analyzing the effects of gene- and genome-level attributes on the frequencies of overlaps.

## INTRODUCTION

Viruses are an enormous part of the natural world, representing the majority of entities in our planet that undergo organic evolution. For instance, a recent study estimated the existence of over 10^31^ bacterial viruses, *i*.*e*, bacteriophage [1], which is only a fraction of viral diversity. A particularly noteworthy feature of virus genomes is the ubiquitous presence of overlapping reading frames (OvRFs): portions of the genome where the same nucleotide sequence encodes more than one protein. OvRFs have been documented in all seven Baltimore classes — categories of viruses by genetic material, including double-stranded DNA (dsDNA) and positive single-stranded RNA (ssRNA+) viruses [2]. A number of hypothetical mechanisms have been proposed to explain this abundance of OvRFs in viruses. First, the prevalence of overlapping genes is hypothesized to be related to genome size. Given that genomes of many viruses are physically constrained by capsid size [3], OvRFs provide a mechanism for encoding more information in a given genome length. Another model proposes that OvRFs could be also used by viruses as a mechanism to accommodate high mutation rates by amplifying the effect size of deleterious mutations (antiredundancy), such that purifying selection removes these mutations more efficiently from the population [4, 5]. In addition, OvRFs have been suggested to be a symptom of gene origination, where a new reading frame may arise within the transcriptional context of an existing reading frame [6]. Recent studies have produced comparative evidence that these *de novo* genes will not initially have a well-established function, but will be able to acquire it over time [7].

Previously, Schlub and Holmes [8] analyzed overlapping genes in 7,450 reference virus genomes in the NCBI viral genomes database [9] to confirm that the number of OvRFs per genome, as well as the number of bases within OvRFs, increases significantly with genome length. In contrast with previous research, however, they also reported that this association was more pronounced in DNA viruses than RNA viruses, and in double-stranded versus single-stranded genomes. Like related work in the literature [3, 5], their comparative study employed quantities like the number of OvRFs or total overlap length (*i*.*e*., the number of nucleotides in overlapping regions) that do not distinguish one reading frame from another. In other words, these are summary statistics where the entire genome is the unit of observation.

Our objective is to incorporate gene homology into characterizing the distribution of OvRFs in virus genomes, with the intent of gaining a more detailed understanding of this phenomenon. This comparative analysis relies on accurate annotation of reading frames in reference genomes. Gene annotation is an increasingly challenging problem, however. For instance, the number of reference virus genomes in the NCBI RefSeq database increased more than five-fold between 2000 and 2015, driven in part by the increasing use of next-generation sequencing platforms [9]. Many putative reading frames in newly discovered virus genomes have no recognizable homologs in protein sequence databases [10]. Furthermore, reading frames in reference genomes are not always annotated with consistent labels, or are assigned the wrong label altogether. Misannotations are sufficiently prevalent that there are multiple collaborations to create and maintain databases of specific categories of genomes with manually-curated gene annotations [11, 12]. To develop a global picture of OvRF diversity across viruses at gene-level resolution, we need an automated method to efficiently label homologous reading frames for related virus genomes.

Here we report a comparative analysis of OvRFs in 12,609 virus reference genomes in the NCBI virus database. First we use conventional genome-level summary statistics to revisit fundamental questions about OvRFs in viruses, *e*.*g*., which frame shifts are most often used by viruses? Next, we develop and employ an alignment-free method for clustering reading frames by sequence homology within a given virus family. This enables us to generate graphs where nodes represent clusters of homologous reading frames. These nodes are connected by two types of edges that indicate the adjacency of reading frames in genomes and the presence of overlaps, respectively. This graph-based approach not only provides an inherent visualization method for the diversity of OvRFs among different virus families, but also enables us to access the rich library of network statistics [13] to characterize the abundance and distribution of OvRFs in virus families.

## MATERIALS AND METHODS

### Data collection and processing

First, we downloaded the accession list of all available virus genomes from the NCBI virus database (https://www.ncbi.nlm.nih.gov/genome/viruses/,accessed on 2020-09-28). This tab-separated file comprised 247,941 rows and six columns labeled as ‘representative’, ‘neighbor’, ‘host’, ‘taxonomy’ and ‘segment name’. Representative genomes are used to denote significant intra-specific variation that cannot be adequately captured by a single reference genome, whereas neighbors are additional validated and complete or nearly-complete genomes for a given species [9]. We used only a single representative genome for each species as sufficient information for our purposes. We used a Python script to retrieve additional metadata (genome length, number of proteins, topology and molecule type) associated with each reference genome using the NCBI Entrez API [14, 15].

The same script was used to generate a tabular dataset recording the genome accession number, product, strand, coordinates and start codon position for every coding sequence (CDS). A second Python script was used to identify putative overlapping reading frames (OvRFs) from the genome coordinates of all CDSs by accession number. Every OvRF was recorded by its location, length in nucleotides, and shift (if applicable) relative to the upstream reading frame. Following convention [16], overlaps between reading frames on the same strand were recorded as +0, +1 and +2 when shifted by zero, one and two nucleotides, respectively. Similarly, overlaps on opposing strands were recorded as −0, −1 and −2 (see Supplementary Figure S1). Next, we extracted Baltimore classifications for virus families from the Swiss-Prot virus annotation resource (https://viralzone.expasy.org [17]).

### Clustering protein data by family

To analyze the distribution of overlapping reading frames in different virus families, we retrieved the protein sequences for all CDSs of all reference genomes of each family from the NCBI virus database. Our objective was to identify homology among protein coding sequences that may be highly divergent and inconsistently annotated at the family level of virus diversity. We also needed to be able to accommodate gene duplication and divergence in DNA viruses, as well as unique reading frames with no homologs in other genomes (*i*.*e*., accessory genes, ORFans [18]). As a result, we decided to use an alignment-free method to compute *k*-mer-based similarity scores between every pair of reading frames within a virus family (Figure S2). We used Python to map each protein sequence to a dictionary of *k*-mer counts for *k* = {1,2,3} as a compact representation of the sparse feature vector. Let *W*(*s*) represent the set of all *k*-mers (words) in a sequence *s*, and let *f* (*s,w*) represent the frequency of *k*-mer *w* in *s*. Using the quantities, we calculated the Bray-Curtis distance [19] between sequences *s* and *t*:

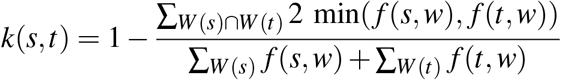

This k-mer distance performed relatively well at the task of protein classification in a recent bench-marking study of alignment-free methods [20], where it was implemented as the intersection distance in the AFKS toolkit [21]. Intuitively, this measure reflects the overlap of two frequency distributions, normalized by the total area of each distribution. The resulting distance matrix was used as input for the t-distributed stochastic neighbor embedding (t-SNE) method implemented in the R package *Rtsne* [22]. This dimensionality reduction method embeds the data points into a lower-dimensional space in such a way that the pairwise distances are preserved as much as possible. Next, we generated a new distance matrix from the coordinates of the embedded points and then used hierarchical clustering using the R function *hclust* with Ward’s criterion [23] (‘ward.D2’). Combining dimensionality reduction and clustering methods is frequently used in combination because distance measures have unexpected properties in high dimensional feature spaces [24].

Finally, we used the R function *cutree* to extract clusters by applying a height cutoff to the dendrogram produced by *hclust*. Increasing the number of clusters by lowering this cutoff accommodates more ORFans. Conversely, raising the cutoff reduces the number of false positive clusters (reading frames that should not be classified as ORFans). To determine an optimal cutoff for a given virus family, we selected the height that balances two quantities. Let *f* (*i, j*) be the number of reading frames assigned to cluster *j* ∈ {1,…, *K*} in genome *i* ∈ {1,…, *N*}. First, we calculated the mean proportion of reading frames with unique cluster assignments per genome:

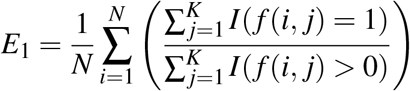

where *I*(*x*) is an indicator function that assumes a value of 1 if *x* is true, and 0 otherwise. Second, we calculated the mean frequency of a cluster assignment across genomes:

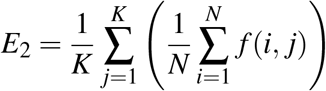

*E*_1_ increases with an increasing number of clusters, whereas *E*_2_ declines because reading frames are distributed across more clusters. Thus, we passed the squared difference (*E*_1_ *− E*_2_)^2^ as an objective function for R function *optimize* to locate the optimal cutoff for each virus family.

### Data visualization

Using the OvRF coordinate data from the preceding analysis, we used a Python script to generate adjacency graphs as node and edge lists for each virus family. Every cluster of homologous reading frames is represented by a node. Each node has two ‘connectors’ representing the 5’ and 3’ ends of the corresponding reading frames in the cluster. For a given genome sequence, all reading frames are sorted by the nucleotide coordinates of their 5’ ends in increasing order. Next, we evaluate every adjacent pair of reading frames in this sorted list. If the 3’ end of the first reading frame occupies a higher coordinate than the 5’ end of another reading frame, then the pair are labelled as overlapping. After screening all adjacent pairs for overlaps, the results were serialized as a weighted graph in the Graphviz DOT language [25], where each node represents a cluster of homologous reading frames. Specifically, we generated two edge lists, one weighted by the frequency that reading frames in either cluster were adjacent in genomes, and a second weighted by the frequency of overlaps. When rendering graphs, we varied edge widths in proportion to the respective weights. In addition, we used the Matplotlib [26] library in Python to visualize the gene order (synteny) of representative genomes in each virus family as concatenated reading frames coloured by cluster assignments, and to visualize the distribution of gene labels by cluster as ‘word clouds’.

### Graph analysis

To analyze the distribution of OvRFs in the context of the adjacency graph of a given virus family, we encoded the numbers of overlaps between every pair of clusters (represented by nodes) as a binomial outcome, given the weight of the corresponding adjacency edge. For every node *A* in the graph, we recorded the number of genomes; number of adjacency edges (degree size); number of triangles (*A* ↔ *B* ↔ *C* ↔ *A*); transitivity (frequency of *B* ↔ *C* given that the graph contains *A* ↔ *B* and *A* ↔ *C*); and the Eigenvector centrality [27], a measure of node importance similar to Google’s PageRank algorithm. Next, we summed these quantities for the two nodes of each edge. We used the resulting values as predictor variables in a zero-inflated binomial regression on the probability of overlap edges, using the *zibinomial* function in the R package VGAM [28]. This mixture model extends the binomial distribution with a third parameter for the probability of zero counts in excess of the binomial. To reduce the chance of overfitting the data, we used stepwise Akaike information criterion (AIC)-based model selection (VGAM function *step4vglm*), where the model search space was limited to the intercept-only model as the lower bound, and the full model with all predictors as the upper bound.

### Code availability

All source code used for our analyses are available under the MIT license at https://github.com/PoonLab/ovrf-viz.

## RESULTS

To examine the distribution of overlapping reading frames (OvRFs) across virus genomes, we retrieved 451,228 coding sequences from 12,609 representative virus genomes as identified by the NCBI virus genomes resource [9]. Based on the annotation of coding sequences (CDS) in each record, we identified 154,687 OvRFs in 6,324 viruses (50.2%). None of the circular ssRNA viral genomes (*n* = 39) contained any OvRFs based on genome annotations. Using the taxonomic annotations in these records, we were able to assign 9,982 (79.2%) of the genomes to Baltimore groups (Figure 1B). Of the remaining 2,627 genomes, 1,001 (38.1%) were comprised of DNA and 1,626 were RNA, based on the molecular type annotation of the respective records.

**Figure 1:**
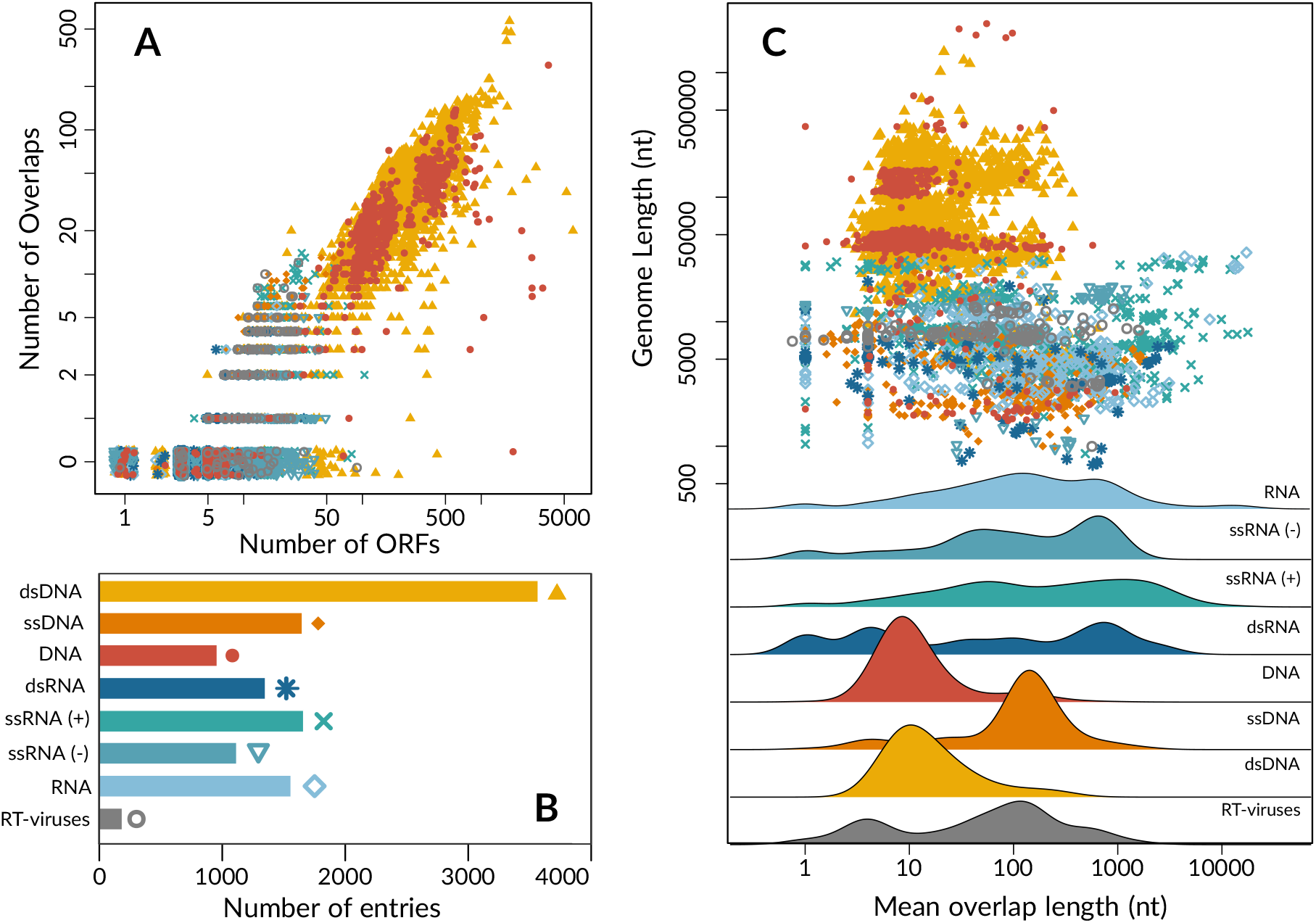
Distribution of overlapping genes across virus genomes. **A**. Scatterplot displaying a positive correlation between the log-transformed numbers of overlapping reading frames (OvRFs) and ORFs per virus genome, stratified by Baltimore class. Genomes with no OvRFs were plotted at 0.5 (labeled ‘0’) with random noise to reduce overplotting. **B**. Barplot of the number of representative virus genomes per Baltimore class, which also serves as a colour and point-type legend for the scatterplots. ‘DNA’ and ‘RNA’ correspond to the molecular type annotations of virus genomes that have not been assigned to a known virus family. **C**. A log-log scatterplot displays the distribution of genomes with respect to overall length (in nucleotides, *y*-axis) and mean length of overlapping regions (*x*-axis) by Baltimore class. Individual plots are provided in Supplementary Figure S3. Underneath, ridgeplots summarize the marginal distributions of genomes with respect to mean overlap lengths, to clarify differences between the Baltimore classes.

### Longer genomes tend to carry shorter overlaps

As expected, the number of OvRFs per genome was positively correlated with the number ORFs (Spearman’s *ρ* = 0.89, *P <* 10^−12^, Figure 1A). In addition, the relative number of OvRFs, *i*.*e*., normalized by the number of ORFs per genome, varied significantly among Baltimore groups (ANOVA, *F* = 835.2, df = 6, *P <* 10^−12^). For instance, double-stranded DNA (dsDNA) viruses — the largest group of viruses in our sample — encode on average 202.8 ORFs and 32.7 overlaps per genome. An extreme case from the Phycodnaviridae family is the Paramecium bursaria Chlorella virus, which infects a eukaryotic algal host and whose genome encodes 1,733 proteins with 541 (31%) overlaps. In contrast, positive sense single-stranded RNA (ssRNA+) viruses encode on average 9.2 ORFs with 2.5 overlaps per genome, and negative sense single-stranded RNA (ssRNA−) viruses encode about 7.1 ORFs and 1.6 overlaps per genome on average. Some RNA virus genomes have abundant overlapping regions, however; *e*.*g*., the simian hemorrhagic fever virus genome (Genbank accession NC 003092) encodes 33 reading frames of which 36% are involved in an overlap.

In contrast, the mean number of nucleotides in overlapping regions was negatively correlated with genome length overall (Spearman’s *ρ* = −0.52, *P <* 10^−12^; Figure 1C). We note that this comparison excludes genomes without any OvRFs, which were significantly shorter (average 6038 nt versus 51424 nt; Wilcoxon rank sum test, *P <* 10^−12^). After adjusting for multiple comparisons (*α* = 6.25 *×* 10^−3^), correlations remained significantly negative within dsDNA, dsRNA, ssRNA+ and unclassified DNA viruses only (Supplementary Figure S3). Correlations within Baltimore classes were largely driven in part by variation among virus families, and we found no consistent trend in correlations within families using a binomial test. While DNA viruses, including single-, double-stranded and unclassified species, tended to carry longer genomes (median 33489 nt, interquartile range (IQR) 2768−59073 nt), their overlapping regions tended to be relatively short (median 15.6 nt, IQR 8.5−61 nt). This trend was largely driven by the dsDNA viruses, and the distributions of overlap numbers and lengths in unclassified DNA genomes (Figure 1C) suggest that these predominantly also represent dsDNA viruses. In comparison, RNA viruses carried fewer but relatively long overlapping regions (median 169.12 nt, IQR 31.34−831.75 nt) for their shorter genome lengths (median 4046 nt, IQR 1986 - 8009 nt).

### Distribution of frameshifts among OvRFs

5,733 (3.7%) of the 154,687 OvRFs identified in our study involved the alternative splicing of one or both transcripts such that there is no consistent relationship between reading frames. These cases are excluded from this section because they complicate the interpretation of frame shifts. The majority (*n* = 92,915, 62.4%) of OvRFs involved reading frames that were shifted by 2 nt on the same strand (+2; Figure 2). These mostly represented dsDNA virus genomes (*n* = 78,191, 84.2%) and comprised almost entirely of overlaps by a single nucleotide (T[AG]ATG) or 4 nt (ATGA). We observed +2 overlaps significantly more often among OvRFs from DNA viruses than from RNA viruses (odds ratio, OR = 3.3; Fisher’s exact test, *P <* 10^−12^). Furthermore, only four out of 29,906 (0.01%) overlaps by 1 nt involved a frame shift other than +2. These four cases involved −2 shifts where one of the reading frames was initiated by the alternate start codon TTG (*e*.*g*., CATTG). Another common type of short OvRF involved −2 frame shifts with an overlap of 4 nt, *e*.*g*., CTAA, where the reverse-complement of TAG is CTA. These were predominantly found in dsDNA virus genomes (*n* = 1423, 73.3%). However, a substantial number (*n* = 419, 21.6%) were also recorded in ssDNA viruses in which a complementary negative-sense strand is generated during virus replication, *e*.*g*., Geminivirus.

**Figure 2:**
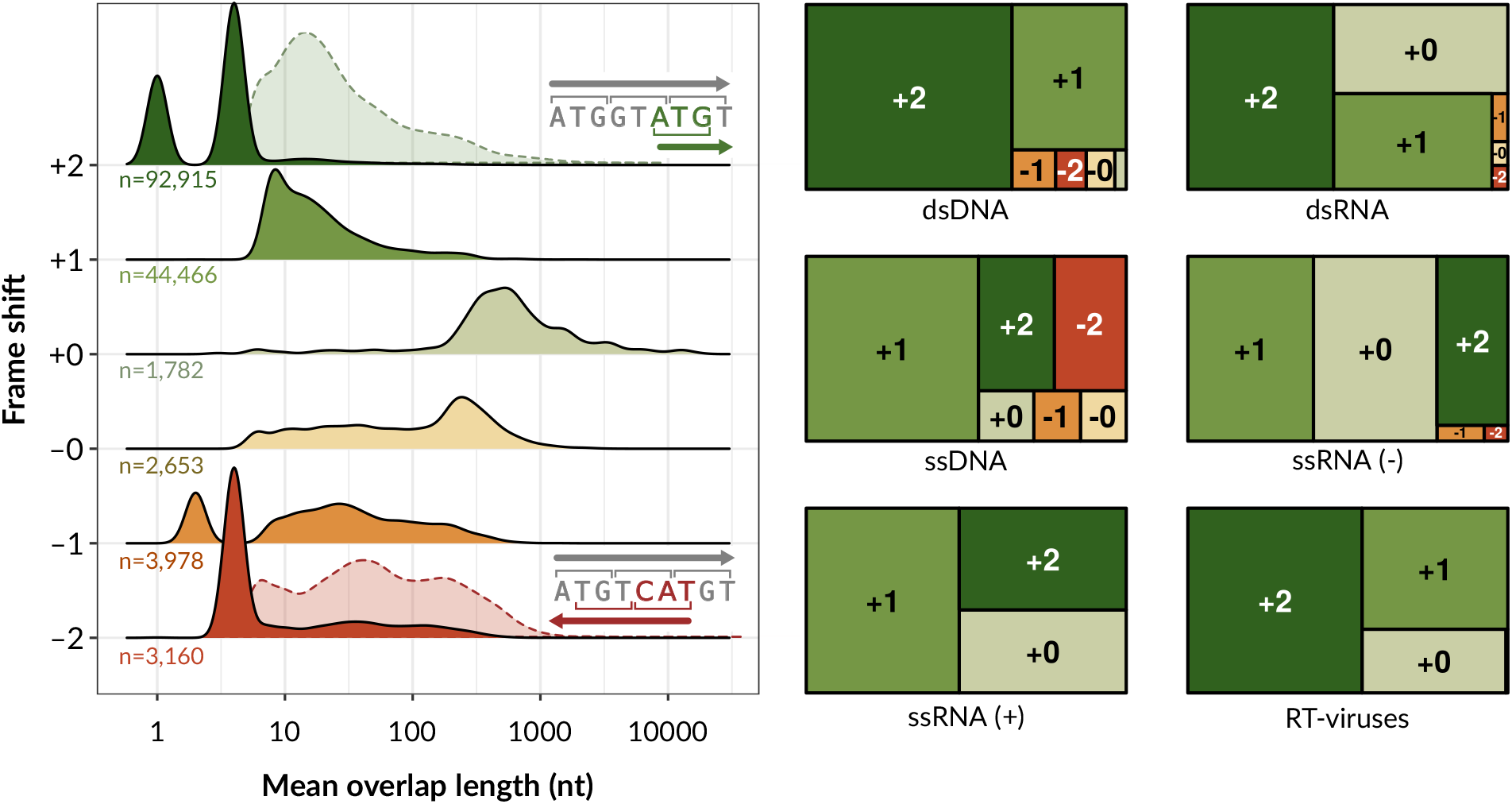
Associations between overlap lengths and frame shifts. (left) Ridgeplots summarizing the distributions of overlap lengths for different frame shifts, where +2 indicates a shift by 2 nt relative to the upstream reading frame, and −2 indicates a 2 nt shift on the opposite strand (note the reverse complement of CAT is ATG). For +2 and −2, we also display the densities after removing overlaps by 1 and 4 nt (dashed outlines), since these predominate the respective distributions. (right) Treemaps summarizing the distribution of frame shifts by Baltimore class. The area of each rectangle is scaled in approximation to the relative frequency of each frame shift.

Excluding OvRFs with short overlaps of 1 or 4 nt, the most common type of OvRF involved a shift of +1. These were observed in both DNA viruses (*n* = 34,175 dsDNA, 1,448 sDNA, and 7,505 unknown) and RNA viruses (*n* = 40 dsRNA, 62 ssRNA−, 917 ssRNA+, and 170 unknown). The median overlap length for +1 OvRFs was 14 nt (IQR 8 to 26 nt). For this type of OvRF, overlaps exceeding 2,000 nt in length were found in ssRNA+ viruses, such as Kennedya yellow mosaic virus (NC 001746) and Providence virus (NC 014126). Overlap lengths tended to be longer for OvRFs with +0 (median 525, IQR 324.8 – 970.5 nt) or −0 (median 114, IQR 27 – 267 nt) frame shifts (Figure 2). We note that OvRFs with a frameshift of +0 tend to represent a reading frame with alternate stop codons, as determined by the number of reading frames with identical start coordinates (*n* = 1349, 75.7%). In addition, 461 of these OvRFs shared the same stop codon; there were only 21 OvRFs with +0 shifts where one reading frame was completely nested within the other.

### Graph-based approach to studying OvRFs

Quantifying OvRFs by statistics like the number of overlaps per genome, or the mean overlap length, reveals substantial variation among Baltimore groups and different frameshifts. However, our objective is to characterize the distribution of OvRFs among virus genomes at a finer resolution. Specifically, these statistics, which are defined at the level of genomes, prevent us from identifying patterns in the distribution of OvRFs at the level of individual genes. For a meaningful comparison at the gene level among virus genomes, we need to be able to identify which genes are homologs. We decided to pursue our objective at the taxonomic level of virus families, to balance diversity in OvRFs with sequence homology. Identifying homologous genes among genomes at the level of virus families is challenging, not only because of substantial evolutionary divergence, but also because genomic rearrangements that can involve the gain, loss or relocation of open reading frames, *i*.*e*., changes in gene order (synteny). For example, the family Rhabdoviridae is characterized for the loss and acquisition of new genes that overlap with consecutive core ORFs, driving substantial variation in genome size and the formation of new accessory genes families [29].

We used an alignment-free *k*-mer-based method [20] to partition all amino acid sequences from genomes in a given virus family into clusters of homology (Supplementary Figure S2). In brief, for each virus family we calculated a *k*-mer distance [19] between every pair of amino acid sequences. We projected the resulting distance matrix into two dimensions by t-distributed stochastic neighbor embedding (t-SNE), and then applied hierarchical clustering to the distances in the 2D plane (Figure 3A). The clustering threshold was determined by balancing the mean frequency of a cluster across genomes against the mean number of unique clusters per genome (Figure 3B).

**Figure 3:**
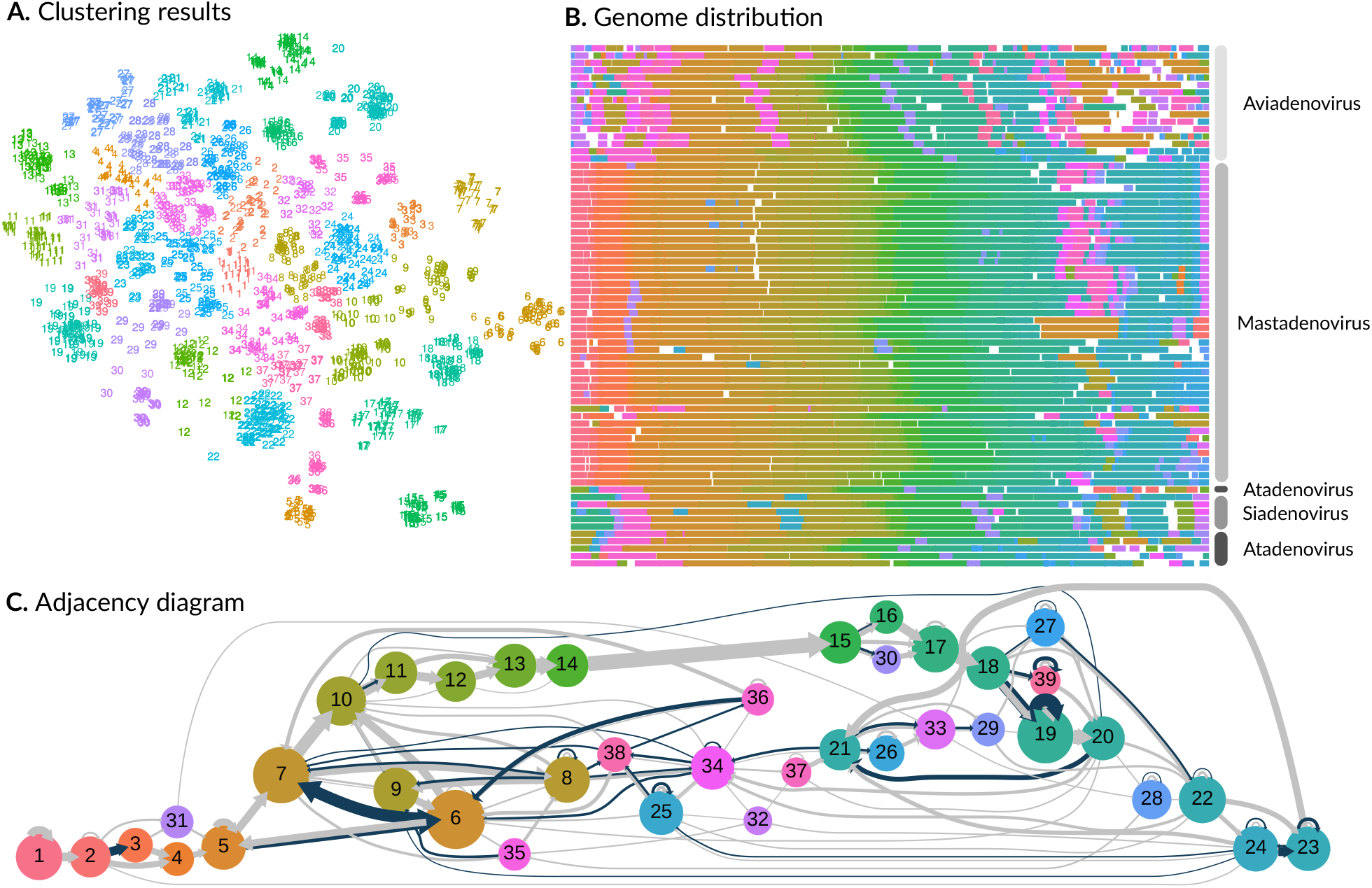
Adenoviridae family analysis. **A**. t-SNE projection of protein sequences from *n* = 71 genomes in the Adenoviridae. Each point represents a protein sequence, coloured and numbered by its cluster assignment. Based on our clustering criteria, we identified a total of 39 clusters for this virus family. **B**. A compact representation of reference genomes labeled by genus. Each set of line segments represent the coding sequences of a genome, coloured by cluster assignments and rescaled to a constant total length. White spaces represent non-coding regions. **C**. A hierarchical layout of the adjacency graph for Adenoviridae. Each node represents a cluster of homologous coding sequences, scaled in proportion to the number of sequences in the cluster. Node numbering and colours were determined by the order of appearance of clusters in the data. Directed edges (arrows) connect nodes representing coding sequences that are adjacent in five or more genomes. Edges are coloured blue if the genes overlap and grey otherwise; widths are scaled in proportion to the number of genomes in either case. This diagram was generated using Graphviz and arrows were manually modified in Inkscape.

We propose a graph-based approach to characterize the distribution of OvRFs in the context of coding sequences in the genome. This approach provides a framework for quantifying overlapping regions at a finer resolution within virus families, and is a natural method for visualizing differences between them. Each node in the graph corresponds to a cluster of homologous coding sequences. Nodes are connected by two sets of directed edges (arrows; Figure 3C). The first set represent the number of genomes in which coding sequences in the respective clusters are located next to each other (adjacency edges). A second set of edges represent the number of genomes in which the adjacent sequences are overlapping (overlap edges). Hence, an overlap edge is never present without an adjacency edge. Because edges are weighted by the number of genomes they each represent, an overlap edge can never have a weight that exceeds the matching adjacency edge.

### Example: Graph-based analysis of Adenoviridae

Adenoviridae is a family of dsDNA viruses with genomes approximately 32,000 nts in length encoding around 30 proteins. Our clustering analysis of protein sequences in the *n* = 72 reference genomes identified 37 clusters (Figure 3A).

Figure 3B displays the distribution of cluster assignments across coding sequences in the genomes. We noted that one of the genomes (bovine adenovirus type 2, NC 002513) had an unusually long non-coding region. We subsequently determined that this reference genome record was not completely annotated, removed it from the dataset and repeated our analysis, resulting in 39 clusters.

The adjacency graph for Adenoviridae features a relatively conserved gene ordering that corresponds to clusters 10 to 20 (Figure 3C). In other words, this part of the graph has a mostly linear structure where nodes tend to have one incoming edge and one outgoing edge. Clusters 10 to 20 correspond to proteins encoded by regions L1-L5 (Supplementary Figure S4). For example, cluster 15 predominately maps to the protein names including the term ‘hexon’. The graph also features several ‘bubbles’ in which one of the coding sequences is gained or lost in a substantial number of genomes. For example, some genomes proceed directly from cluster 11 (pVII) to 13 (pX), by-passing 12 (pV). Similarly, cluster 31 (IX or ORF0) is gained in at least 11 genomes. In addition, the graph splits between clusters 19 and 39 as it traverses from cluster 18 to 20. These clusters correspond to mixtures of the proteins 33K and 22K, which are both encoded by alternative splices of the same gene transcript that includes a long intron [30]. This split, as well as self-loop edges on both clusters, implies that our clustering method can be confounded by inconsistent annotation of such isoforms.

The graph also contains distinct groups of nodes with multiple incoming or outgoing edges, which represent homologous clusters of coding sequences with more variable orderings in Adenoviridae. For example, clusters 8, 34, 36 and 38 generally correspond to reading frames in the E1 region of aviadenovirus (genus of bird-associated adenovirus) genomes associated with gene duplication events [31]. Similarly, cluster 25 maps to the RH family of duplicated genes in aviadenovirus and atadenovirus genomes. The presence of homologous coding sequences with potentially common origins in both the E1 (5’) and E4 (3’) regions of the Adenoviridae genomes induces the overall cyclic structure in this adjacency graph.

In Adenoviridae, OvRFs vary from 1 to 19 nt in length, with a median length of 10 nt (IQR 8 – 12 nt). By visualizing clusters of homologous coding sequences as a graph, we can see that the conserved ‘backbone’ of L1-L5 genes are relatively free of overlaps. In addition, Figure 4 displays the adjacency graphs produced for four other virus families (Rhabodiviridae, Geminiviridae, Coronaviridae and Papillomaviridae, representing four different Baltimore groups. This visual comparison of adjacency graphs not only clarifies the substantial variation in the frequency of OvRFs among families, but also reveals differences in the distribution of OvRFs among reading frames. For example, OvRFs in genomes of Geminiviridae tended to be associated with common adjacency edges on the ‘left’ side of the graph, corresponding to homologous reading frames closer to the 5’ end of genomes. OvRFs in Coronaviridae genomes tended to be associated with less common pairs of adjacent reading frames (*e*.*g*., 10-7 and 7-3 versus 8-4 and 4-5; Figure 4).

**Figure 4:**
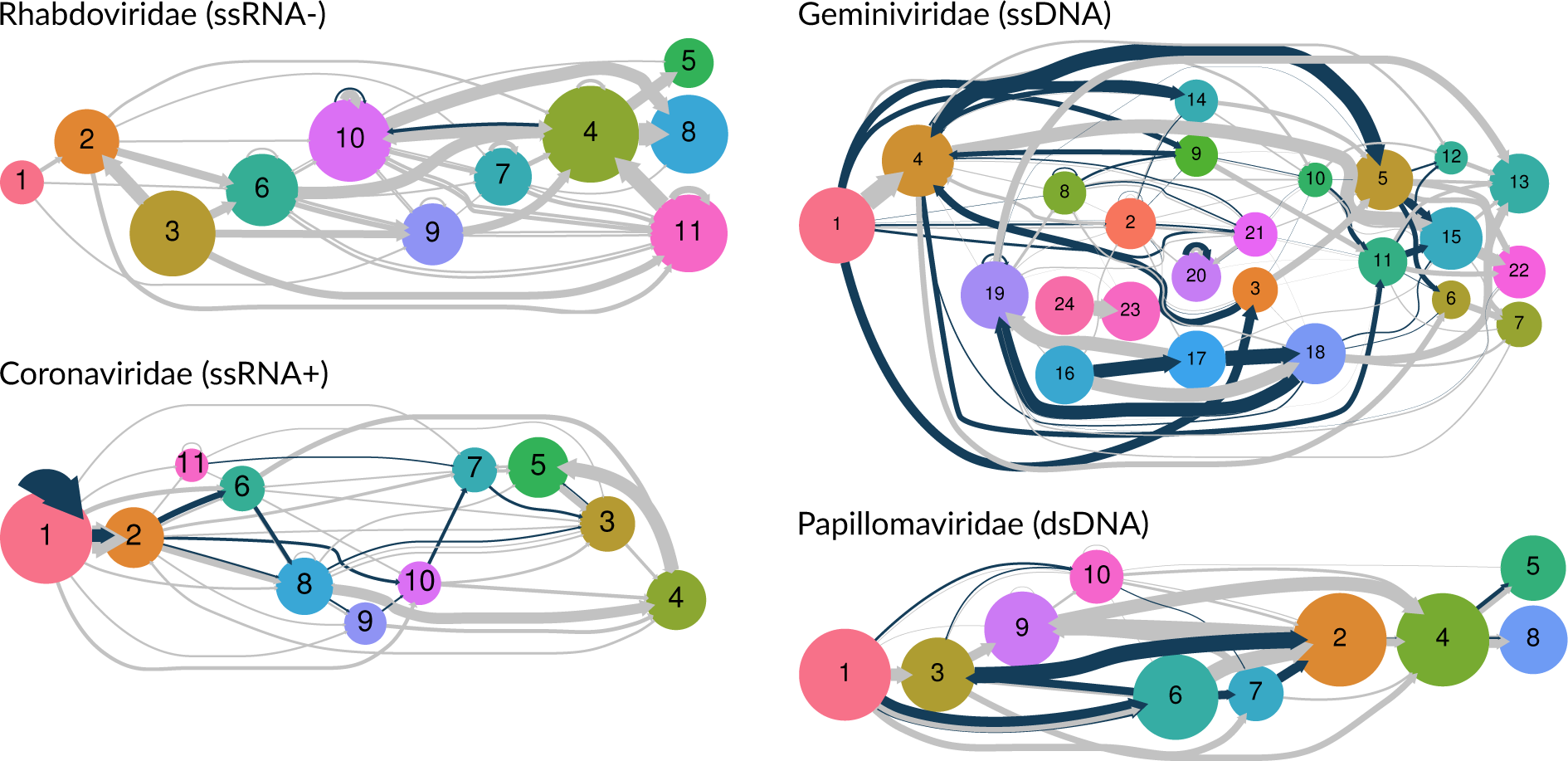
Adjacency graphs for different virus families. These graphs were generated from the clustered reading frame data, using the same procedure that we employed to generate Figure 3 for Adenoviridiae. Blue edges indicate overlapping reading frames, and grey edges represent reading frames that are adjacent but not overlapping. Edge widths were rescaled by a factor of 0.5 for Geminiviridae and Papillomaviridae to accommodate differences in sample size (numbers of genomes) among virus families. Arrowheads were manually adjusted in Inkscape as in Figure 3.

### Variation among families

To analyze the distributions of OvRFs in the context of adjacency graphs, we fit a zero-inflated binomial regression model to the weights of overlap and adjacency edges for every pair of clusters. For example, out of 69 genomes with adjacent coding sequences assigned to clusters 6 and 7 in the Adenoviridae graph, 57 genomes had an overlap between the sequences and 12 did not. We calculated the number of genomes, degree size, number of triangles, transitivity and Eigenvector centrality as edge-level attributes, and used a stepwise model selection procedure to determine which combination of attributes was best supported by the data as predictor variables. The results of fitting these regression models to each graph are summarized in Figure 5. Effect size estimates varied substantially among virus families. For example, centrality was significantly associated with higher probabilities of overlapping reading frames for Geminiviridae, Papillomaviridae and Rhabdoviridae, but with lower probabilities in Coronaviridae. A cluster with high centrality is connected to many other clusters that are also of high degree size. In our context, high degree sizes correspond to reading frames with variable neighbours or diverse locations in a genome, *e*.*g*., due to multiple gene capture and duplication events [31]. Triangles in the adjacency graph tended to be associated with lower rates of overlap. For instance, gene loss by deletion (from *A → B → C* to *A → C*) is more likely to be tolerated if the adjacent reading frames *A* and *C* do not overlap with the targeted gene *B*. For Adenoviridae, transitivity had a relatively slight but significant negative effect on overlap probability (Figure 5). This is consistent with our previous observation that clusters comprising a core ‘backbone’ in the adjacency graph tended to be associated with fewer overlap edges.

**Figure 5:**
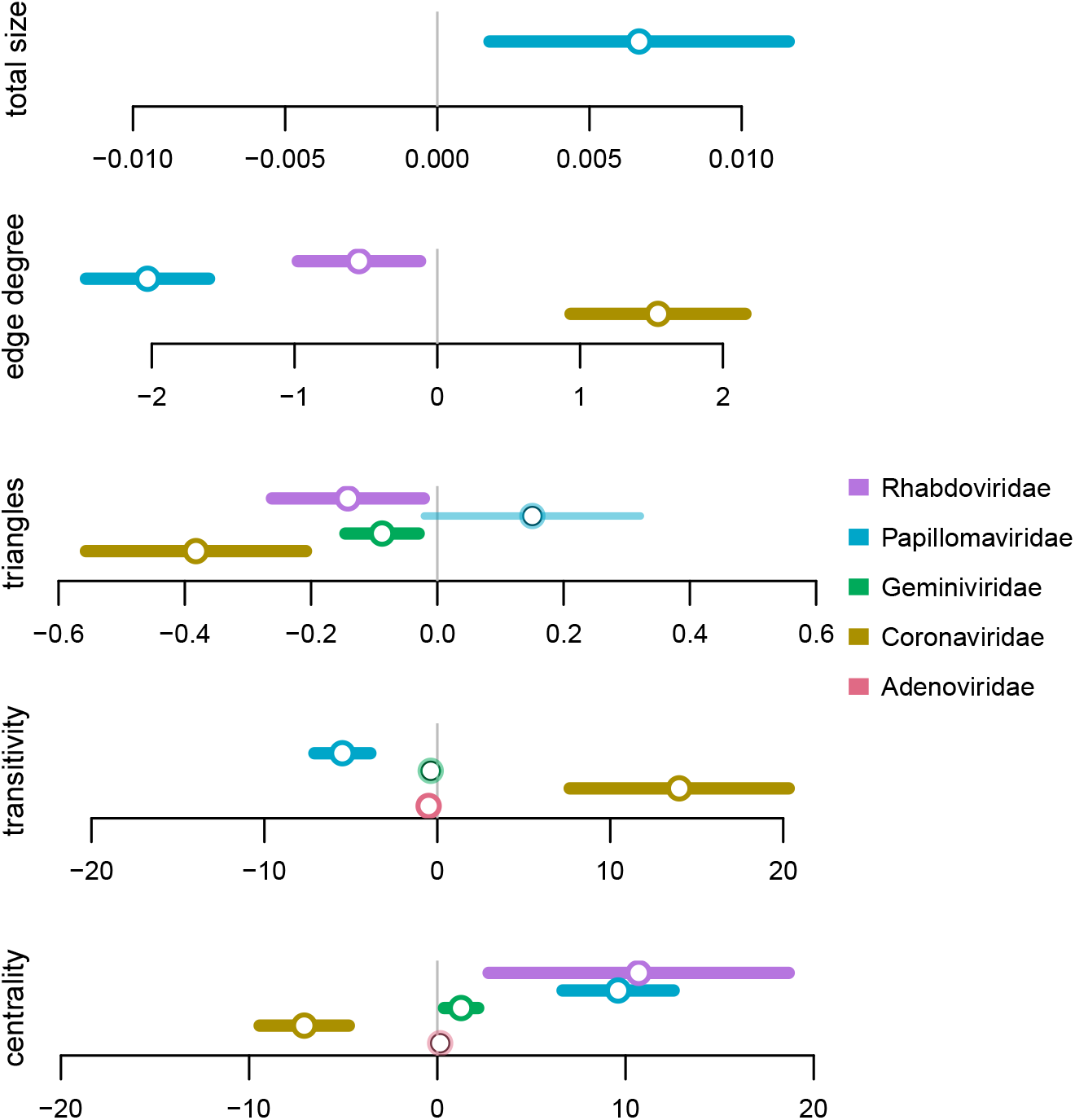
Forest plot of zero-inflated binomial regressions on adjacency graphs. Points and lines are omitted for terms that were discarded by a stepwise AIC model selection procedure. Each point corresponds to coefficient estimates from zero-inflated binomial regressions on the probability of an overlap between adjacent reading frames, for clustered reading frames from the genomes of each of five virus families (see colour legend). Line segments correspond to the 95% confidence interval of the estimate, drawn in bold when the interval does not include zero. Total size = total number of genomes with adjacent reading frames assigned to the respective clusters. Edge degree = total edge degree of the linked clusters. Triangles = total number of triangles involving either cluster. Transitivity = total transitivity of the linked clusters. Centrality = total Eigenvector centrality of the linked clusters.

## DISCUSSION

The number of genome sequences for different viruses in public databases is rapidly growing, driven in part by environmental metagenomic sequencing projects [32] and education/outreach programs like SEA-PHAGES [33]. These data provide a significant opportunity to examine the composition of these genomes to identify large-scale patterns. Here we have characterized the distribution of overlapping reading frames (OvRFs) in over 12,600 annotated virus genomes. Beginning with conventional comparative methods, we first confirmed previous findings that overlapping genes are ubiquitous across all Baltimore classes, with examples identified in 50.2% of the virus genomes. We observed that the majority of non-splicing OvRFs are short (*e*.*g*., less than 10 nucleotides). However, the small overlaps in our study were predominantly by 1 or 4 nucleotides, whereas previous work [8] reported peaks at slightly longer lengths (3 and 7 nucleotides, respectively). We also confirmed previous reports [2, 3, 5] that the number of OvRFs increases with genome length, whereas OvRFs tend to be shorter in longer genomes. These trends are consistent with the compression theory that proposes that overlapping genes are a significant mechanism for reducing genome lengths [3]. However, we must be cautious about interpreting these patterns because, like previous work, there is no adjustment for non-independence among observations due to evolutionary homology, *i*.*e*., identity by descent. This can be mitigated in part by examining correlations within virus families. At this level, we did not find significant evidence of a consistent association between overlap and genome lengths (Supplementary Figure S3).

In the absence of a standard notation for frameshifts in overlapping reading frames, past studies have devised different labeling systems (Supplementary Figure S1). For overlapping reading frames on the same strand, we labeled the frameshift relative to the ‘upstream’ reading frame. Following [16], we used a negative sign to indicate that overlapping reading frames are on opposite strands. However, we used −1 and −2 to indicate that the codons on the opposite strand are shifted by one and two nucleotides, respectively, relative to the −0 frame, which we consider to be a more intuitive notation. In an analysis of 701 RNA virus genomes, Belshaw and collaborators [3] previously reported that most overlapping genes consist of +1 and +2 frameshifts (+1 and −1 in their notation). We observed similar results in our analysis of 5,972 RNA virus genomes, which we further stratified by Baltimore class (Figure 2). Note that, unlike some earlier work, we included +0 overlaps in our database. Even though the reading frames share codons, they yield different gene products such that the codons are exposed to different selective environments. These cases may thereby provide a means of differentiating between the compression [3] and antiredundancy [4] hypotheses of OvRFs in viruses, since +0 overlaps increase the selective burden of the same nucleotides, whereas other frameshifts increase the number of nucleotide sites under purifying selection. This is countered by a narrower repertoire of protein sequences and structures.

In our data, antisense frameshifts (*i*.*e*., −0, −1 and −2) account for only the 6.6% of all overlaps and are primarily found in DNA virus genomes. We detected a total of 14 cases of antisense frameshifts in RNA virus genomes (two instances in −2, four in −1, and eight in −0). For example, two −1 overlaps 434 and 44 nt in length are annotated in segment S of the dsRNA virus Pseudomonas phage phiYY (NC 042073). Since these involve hypothetical proteins in a recently discovered dsRNA phage, however, we must be cautious about interpreting these results. In DNA viruses, −1 are slightly more common than −0 and −2 OvRFs (Figure 2), especially if we exclude the most common −2 overlap by four nucleotides (*i*.*e*., CTAA). However, Lébre and Gascuel [16] recently determined that the −1 frameshift (−2 in their notation) was the most constrained, in that the codons used in one reading frame limit the amino acids that can be encoded in the other. It also minimizes the expected frequency of stop codons in the opposing strand, but −1 overlaps were not significantly longer than other types in our data.

One of the key challenges for extending our comparative analysis to the level of individual reading frames was the assignment of reading frames into clusters of homology. This is complicated not only by extensive sequence divergence at the level of virus families, but also the gain or loss of reading frames in different lineages through processes that include gene duplication. Furthermore, the annotation of reading frames in a general purpose public database like Genbank is not sufficiently consistent to rely on these labels. For example, cluster 15 in our analysis of Adenoviridae genomes comprised coding sequences with diverse labels, including ‘hexon’, ‘hexon protein’, ‘hexon capsid protein’, ‘L3 hexon’, ‘II’, ‘capsid protein II’, ‘protein II’, and the ubiquitous ‘hypothetical protein’ label (Supplementary Figure S4). We also found several examples of genomes in the NCBI virus database in which reading frames were incompletely annotated. The bovine adenovirus type 2 reference genome (NC 002513), for example, has only 11 annotated coding sequences. Adenovirus genomes typically contain about 30 to 40 genes. Since this reference genome lacks coding sequence annotations over a 15 kbp interval, it was apparent that many genes were simply not annotated. We subsequently confirmed this using a gene prediction and homology search analysis and discarded this reference genome from our analysis.

Given the abundance of genomic diversity at the level of virus families, the assignment of reading frames into homologous groups is not unambiguous. Thus, we utilized an alignment-free approach to cluster the coding sequences in the reference genomes for each virus family. There are a large number of alignment-free methods that extract k-mers from two input sequences (see [20] for a recent review). We chose the Bray-Curtis distance (also known as the intersection distance) because it performed comparatively well at the task of protein classification in a recent bench-marking study [20]. However, the classification analysis in that study was performed on protein databases curated to span a broad range of relationships, including both cases of evolutionary and structural homology. In addition, it was necessary to apply a threshold to our hierarchical clustering results to extract distinct clusters. At any threshold, it is inevitable that some number of reading frames were misclassified into separate clusters (false negatives) or into the same cluster (false positives). For each virus family, we selected thresholds that minimized the number of duplicate cluster assignments per genome while maximizing the overall frequencies of clusters across genomes. Although it is intuitive, we must acknowledge that this is an *ad hoc* approach to the hierarchical clustering of grouped data without a known theoretical basis. However, we contend that such clustering algorithms will become increasingly important with a rapidly growing number of new virus genomes from the metagenomic analysis of environmental samples.

To evaluate the role of OvRFs in the emergence of genes *de novo* by overprinting [5, 6], we need to characterize the distribution of overlaps at the level of individual reading frames. We employed unsupervised clustering of homologous reading frames to identify gene order polymorphisms at the level of virus families. Visualizing the syntenic relationships among clusters at this taxonomic level provided some interesting patterns. For example, members of Adenoviridae have about 16 conserved ‘core’ genes in the middle of the genome that are responsible for DNA replication and encapsidation, and the formation and structure of the virion [31]. These core genes formed a distinct backbone in our adjacency graph of this family that was relatively free of overlaps (Figure 3). On the other hand, cluster assignments tended to become more variable towards the 5’ and 3’ ends of the genome in association with an increasing frequency of OvRFs. Similarly, OvRFs in Coronaviridae tended to be associated with pairs of clusters representing accessory genes that were less frequently adjacent in these genomes (Figure 4). As a member of the virus order nidovirales, coronaviruses have undergone extensive selection to expand the repertoire of genes encoded by their relatively long RNA genomes [34]. Accessory genes assigned to clusters with overlaps tended to be found in varying locations in genomes of Coronaviridae, which is more consistent with gene duplication and insertion than overprinting.

In summation, we have described and demonstrated a new approach to characterize the distribution of overlapping reading frames in diverse virus genomes. Adjacency graphs provide a framework for both visualizing these distributions and for hypothesis testing, *i*.*e*., effects of gene- or genome-level attributes on the frequencies of overlaps between specific reading frames. In future work, we will develop comparative methods on the topologies and features of adjacency graphs to identify shared characteristics between virus families at this level. We further postulate that adjacency graphs may provide useful material for extending methods for ancestral gene order reconstruction [35], where the graphs can address the problem of uncertain labelling of genes. Ideally, one would simultaneously reconstruct the phylogeny relating observed genomes. Reconstructing ancestral gene order is already an NP-hard problem [36]. Given the diverse and evolutionarily fluid composition of many virus genomes, however, it is remarkable that the gain and loss of reading frames has not been explored as much as larger organismal genomes.

## ACKNOWLEDGEMENTS

This work was supported by a Discovery Grant from the Natural Sciences and Engineering Research Council of Canada (05516-2018 RGPIN) to AFYP.

## SUPPLEMENTARY FIGURES

**Figure S1:**
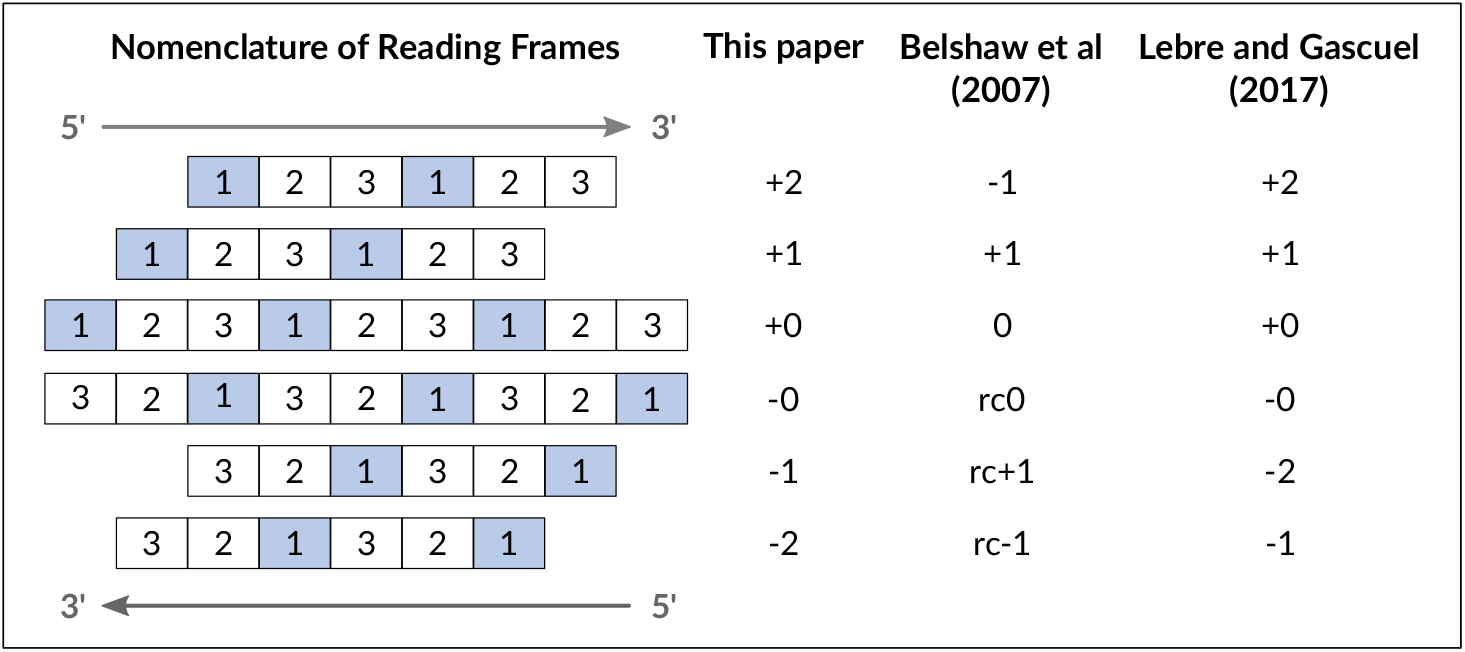
Notation of the 6 possible frame shifts used in this study and two other studies [3, 16].

**Figure S2:**
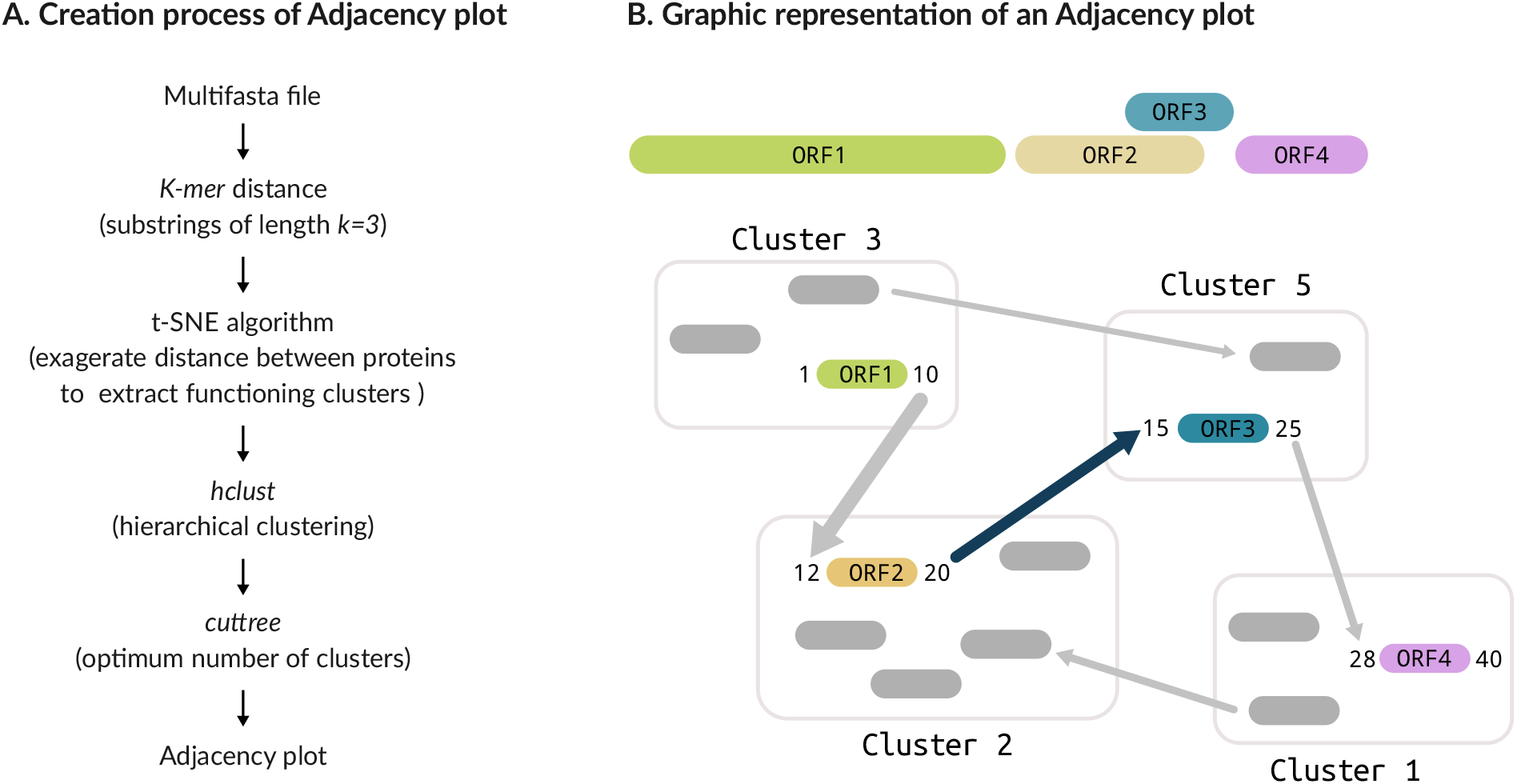
Creation of adjacency plots. **A**. Steps used to generate the input file for the adjacency plot. First, we downloaded a multifasta file containing the protein sequences of the reference genomes for each species in the virus family. Then, we used a Python script to calculate the *k-mer* distance between proteins followed by an R script to designate each protein to a cluster according to homology. Finally, we used a Python script to generate dot files using Graphviz. **B**. Adjacency plot interpretation. Each one of the proteins that constitute a genome is assigned to a different cluster with homologous proteins from other species. From each cluster, we draw arrows in gray that represent adjacent proteins and arrows in blue that represent overlapping proteins. The width of the arrow is proportional to the number of proteins related between the two clusters. One cluster can have entries adjacent to proteins in different clusters. In this example, cluster 3 has proteins adjacent to proteins in cluster 2 and cluster 5.

**Figure S3:**
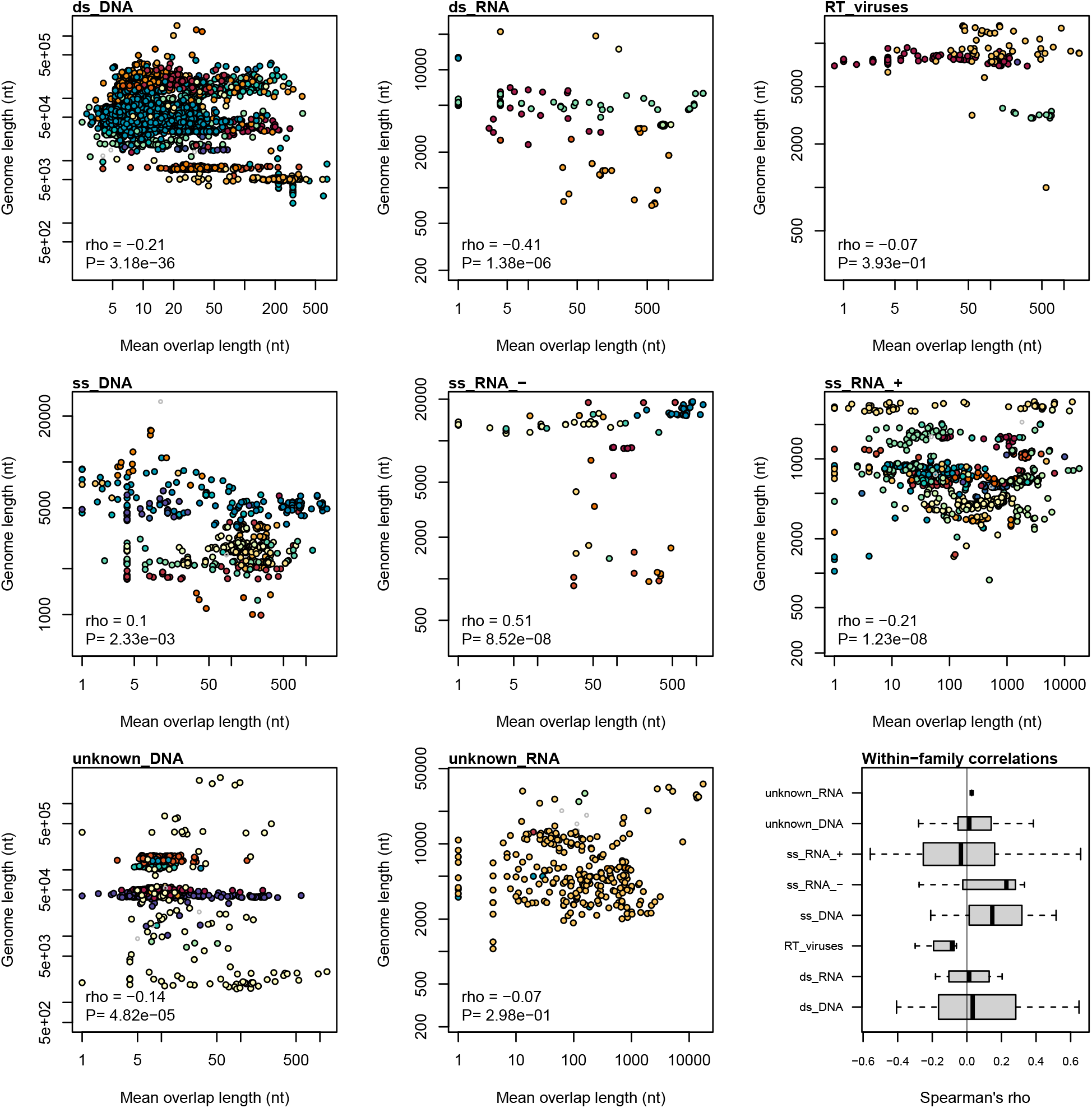
Scatterplots of genome length on mean overlap length by Baltimore class. Each point represents a virus genome, coloured by virus family. Families represented by only one genome were coloured in grey (smaller point size). Results from Spearman rank correlations are summarized in the lower left corner of each plot. The lower-right panel displays a box- and-whisker plot summarizing the distributions of Spearman’s rank correlation coefficients (*ρ*) for genomes within families, grouped by Baltimore class.

**Figure S4:**
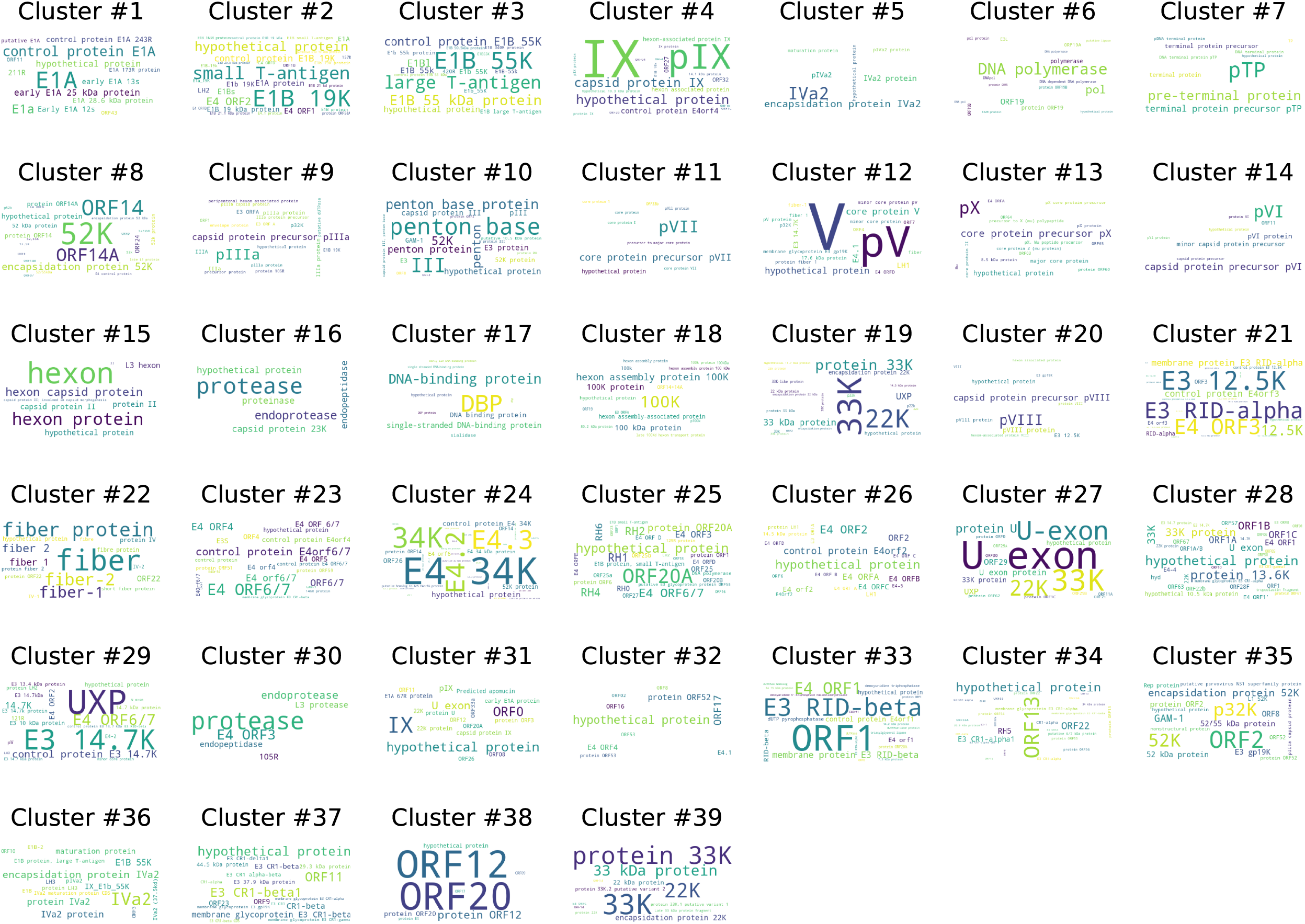
Word clouds of protein names mapped to clusters for Adenoviridae.

## REFERENCES

1. Cobián Güemes AG, Youle M, Cantú VA, Felts B, Nulton J, Rohwer F. Viruses as winners in the game of life. Annual Review of Virology. 2016;3:197–214.

2. Brandes N, Linial M. Gene overlapping and size constraints in the viral world. Biology direct. 2016;11(1):26.

3. Belshaw R, Pybus OG, Rambaut A. The evolution of genome compression and genomic novelty in RNA viruses. Genome research. 2007;17(10):1496–1504.

4. Krakauer DC, Plotkin JB. Redundancy, antiredundancy, and the robustness of genomes. Proceedings of the National Academy of Sciences. 2002;99(3):1405–1409.

5. Chirico N, Vianelli A, Belshaw R. Why genes overlap in viruses. Proceedings of the Royal Society B: Biological Sciences. 2010;277(1701):3809–3817.

6. Sabath N, Wagner A, Karlin D. Evolution of viral proteins originated de novo by overprinting. Molecular biology and evolution. 2012;29(12):3767–3780.

7. Willis S, Masel J. Gene birth contributes to structural disorder encoded by overlapping genes. Genetics. 2018;210(1):303–313.

8. Schlub TE, Holmes EC. Properties and abundance of overlapping genes in viruses. Virus evolution. 2020;6(1):veaa009.

9. Brister JR, Ako-Adjei D, Bao Y, Blinkova O. NCBI viral genomes resource. Nucleic acids research. 2015;43(D1):D571–D577.

10. Dutilh BE, Cassman N, McNair K, Sanchez SE, Silva GG, Boling L, et al. A highly abundant bacteriophage discovered in the unknown sequences of human faecal metagenomes. Nature communications. 2014;5(1):1–11.

11. Bernt M, Donath A, Jühling F, Externbrink F, Florentz C, Fritzsch G, et al. MITOS: improved de novo metazoan mitochondrial genome annotation. Molecular phylogenetics and evolution. 2013;69(2):313–319.

12. Grazziotin AL, Koonin EV, Kristensen DM. Prokaryotic Virus Orthologous Groups (pVOGs): a resource for comparative genomics and protein family annotation. Nucleic acids research. 2016;p. gkw975.

13. Newman M. Networks. Oxford University Press; 2018.

14. Cock PJ, Antao T, Chang JT, Chapman BA, Cox CJ, Dalke A, et al. Biopython: freely available Python tools for computational molecular biology and bioinformatics. Bioinformatics. 2009;25(11):1422–1423.

15. Database resources of the national center for biotechnology information. Nucleic acids research. 2018;46(D1):D8–D13.

16. Lebre S, Gascuel O. The combinatorics of overlapping genes. Journal of theoretical biology. 2017;415:90–101.

17. Hulo C, De Castro E, Masson P, Bougueleret L, Bairoch A, Xenarios I, et al. ViralZone: a knowledge resource to understand virus diversity. Nucleic acids research. 2011;39(suppl 1):D576–D582.

18. Yin Y, Fischer D. Identification and investigation of ORFans in the viral world. Bmc Genomics. 2008;9(1):1–10.

19. Bray JR, Curtis JT. An ordination of the upland forest communities of southern Wisconsin. Ecological monographs. 1957;27(4):325–349.

20. Zielezinski A, Girgis HZ, Bernard G, Leimeister CA, Tang K, Dencker T, et al. Benchmarking of alignment-free sequence comparison methods. Genome biology. 2019;20(1):144.

21. Luczak BB, James BT, Girgis HZ. A survey and evaluations of histogram-based statistics in alignment-free sequence comparison. Briefings in bioinformatics. 2019;20(4):1222–1237.

22. Van Der Maaten L. Accelerating t-SNE using tree-based algorithms. The Journal of Machine Learning Research. 2014;15(1):3221–3245.

23. Murtagh F, Legendre P. Wards hierarchical agglomerative clustering method: which algorithms implement Wards criterion? Journal of classification. 2014;31(3):274–295.

24. Ronan T, Qi Z, Naegle KM. Avoiding common pitfalls when clustering biological data. Science signaling. 2016;9(432):re6–re6.

25. Gansner E, Koutsofios E, North S. Drawing graphs with dot. Technical report, AT&T Research. http://www.graphviz.org/Documentation; 2006.

26. Hunter JD. Matplotlib: A 2D graphics environment. Computing in Science & Engineering. 2007;9(3):90–95.

27. Ruhnau B. Eigenvector-centralitya node-centrality? Social networks. 2000;22(4):357–365.

28. Yee TW. The VGAM package for categorical data analysis. Journal of Statistical Software. 2010;32(10):1–34.

29. Walker PJ, Firth C, Widen SG, Blasdell KR, Guzman H, Wood TG, et al. Evolution of genome size and complexity in the Rhabdoviridae. PLoS Pathog. 2015;11(2):e1004664.

30. Reddy PS, Idamakanti N, Zakhartchouk AN, Baxi MK, Lee JB, Pyne C, et al. Nucleotide sequence, genome organization, and transcription map of bovine adenovirus type 3. Journal of virology. 1998;72(2):1394–1402.

31. Davison AJ, Benko? M, Harrach B. Genetic content and evolution of adenoviruses. Journal of General Virology. 2003;84(11):2895–2908.

32. Mokili JL, Rohwer F, Dutilh BE. Metagenomics and future perspectives in virus discovery. Current opinion in virology. 2012;2(1):63–77.

33. Jordan TC, Burnett SH, Carson S, Caruso SM, Clase K, DeJong RJ, et al. A broadly implementable research course in phage discovery and genomics for first-year undergraduate students. MBio. 2014;5(1).

34. Gorbalenya AE, Enjuanes L, Ziebuhr J, Snijder EJ. Nidovirales: evolving the largest RNA virus genome. Virus research. 2006;117(1):17–37.

35. Yang N, Hu F, Zhou L, Tang J. Reconstruction of ancestral gene orders using probabilistic and gene encoding approaches. PloS one. 2014;9(10):e108796.

36. Figeac M, Varré JS. Sorting by reversals with common intervals. In: International Workshop on Algorithms in Bioinformatics. Springer; 2004. p. 26–37.

